# The structure of *Vibrio cholerae* FeoC reveals conservation of the helix-turn-helix motif but not the cluster-binding domain

**DOI:** 10.1101/2022.02.26.482101

**Authors:** Janae B. Brown, Mark A. Lee, Aaron T. Smith

## Abstract

Most pathogenic bacteria require ferrous iron (Fe^2+^) in order to sustain infection within hosts. The ferrous iron transport (Feo) system is the most highly-conserved prokaryotic transporter of Fe^2+^, but its mechanism remains to be fully characterized. Most Feo systems are composed of two proteins: FeoA, a soluble SH3-like accessory protein and FeoB, a membrane protein that translocates Fe^2+^ across a lipid bilayer. Some bacterial *feo* operons encode FeoC, a third soluble, winged-helix protein that remains enigmatic in function. We previously demonstrated that select FeoC proteins bind O_2_-sensitive [4Fe-4S] clusters via Cys residues, leading to the proposal that some FeoCs could sense O_2_ to regulate Fe^2+^ transport. However, not all FeoCs conserve these Cys residues, and FeoC from the causative agent of cholera (*Vibrio cholerae*) notably lacks any Cys residues, precluding cluster binding. In this work, we determined the NMR structure of *Vc*FeoC, which is monomeric and conserves the helix-turn-helix domain seen in other FeoCs. In contrast, however, the structure of *Vc*FeoC reveals a truncated winged β-sheet in which the cluster-binding domain is notably absent. To test the interactions of *Vc*FeoC with *Vc*FeoB, we used NMR to demonstrate that these proteins interact in a 1:1 stoichiometry with a K_d_ of *ca*. 25 μM. Finally, using homology modeling, we predicted the structure of *Vc*NFeoB and used docking to identify an interaction site with *Vc*FeoC, which is confirmed by NMR spectroscopy. These findings provide the first atomic-level structure of *Vc*FeoC and contribute to a better understanding of its role vis-à-vis FeoB.

## Introduction

Iron is essential for nearly all organisms, as it is required for indispensable cellular processes from electron transport and ATP synthesis to DNA biosynthesis [1, 2]. Given this importance, the acquisition of iron is thus necessary for the survival of nearly every organism. For many pathogenic bacteria, iron is typically obtained from a host as siderophore-bound ferric iron (Fe^3+^), iron protoporphyrin IX (heme), and/or ferrous iron (Fe^2+^), and the acquisition of this element is necessary to establish and to maintain infection [2–7]. Methods of Fe^3+^ and heme acquisition have been well-characterized, but pathways for Fe^2+^ uptake are less well-understood.

The ferrous iron transport system (Feo) is the most conserved and broadly-distributed system dedicated to Fe^2+^ transport in prokaryotes [5]; however, the precise mechanism of Feo-mediated iron transport remains unclear. The *feo* operon is generally bipartite and encodes for FeoA, a small (*ca*. 8 kDa), cytosolic SH3-like protein [8] and for FeoB, a large (*ca*. 85 kDa) transmembrane protein capable of NTP hydrolysis at its soluble N-terminal domain (typically termed NFeoB) [9]. However, in approximately 13% of bacteria the *feo* operon is tripartite and additionally encodes for FeoC, a small (*ca*. 9 kDa), cytosolic protein [10–12]. Structures of FeoC have demonstrated its architecture to include a conserved, trihelical helix-turn-helix (HTH) domain fused to a winged β-sheet, akin to that of the LysR transcriptional regulator (LTTR) family [10, 11, 13]. This structural similarity has led to proposals that FeoC may have a function in transcriptional regulation [14], but this hypothesis has not been verified experimentally. Additionally, sequence alignments of FeoC proteins highlight the strong conservation of Cys residues within the winged β-sheet, which initially suggested an iron-dependent function of FeoC that could be similar to the iron-sensing diphtheria toxin repressor (DtxR) [15, 16].

Our lab recently determined that *Escherichia coli* and *Klebsiella pneumoniae* FeoCs (*Ec* and *Kp*FeoC, respectively) bind [4Fe-4S] clusters using their Cys-rich winged β-sheet [16, 17]. Although the specific impact of cluster binding on iron transport is currently unknown, we demonstrated that this cluster binding event induces conformational changes in FeoC, which we posited could trigger FeoC-mediated regulation of Feo function, perhaps through interactions with FeoB at its cytosolic domain [17]. Notably, an X-ray crystal structure of the N-terminal domain of FeoB (NFeoB) from *K. pneumoniae* in complex with *Kp*FeoC has been determined (PDB ID 4AWX; [10]) but the winged β-sheet including its [Fe-S] cluster-binding domain was disordered, precluding assignments of protein-protein interactions of this domain. We also demonstrated that the [4Fe-4S] cluster rapidly degrades upon O_2_ exposure, which led to the hypothesis that FeoC may regulate Feo function by sensing O_2_ at the [Fe-S] cluster, similar to the fumarate and nitrate reductase (FNR) response regulator [17–19]. Unfortunately, this rapid reactivity in the presence of minute amounts of O_2_ made characterizing the structure of the [4Fe-4S] cluster-bound form of FeoC difficult even under anoxic conditions. However, some FeoC proteins, including *Vibrio cholerae* FeoC (*Vc*FeoC), are required for iron transport but do not feature the Cys residues necessary for [Fe-S] cluster-binding based on sequence predictions [12, 20, 21]. Thus, we propose that *Vc*FeoC may belong to a class of FeoC proteins that do not require [Fe-S] cluster-binding and may not be iron-regulated directly but could maintain a state of constitutive activity in their interactions with FeoB [12].

To this end, we employed solution NMR spectroscopy to determine the first threedimensional structure of *Vc*FeoC. Gel filtration and NMR data demonstrate that *Vc*FeoC is monomeric in solution under the conditions employed. Importantly, our new structure shows that *Vc*FeoC bears a HTH domain conserved among FeoCs; however, the winged β-sheet is shortened and compacted relative to other FeoC proteins and does not conserve the [Fe-S] cluster-binding domain. To test whether *Vc*FeoC could bind to *Vc*FeoB in the absence of metal, we orthogonally cloned, expressed, solubilized, and purified intact *Vc*FeoB and performed 2D NMR titration assays. These studies allowed us to approximate the binding between *Vc*FeoB and *Vc*FeoC, and to map *Vc*FeoC residues that contribute to the binding interface. Finally, we generated a homology model of the soluble N-terminal domain of *Vc*FeoB (*Vc*NFeoB) and performed docking studies in an effort to identify regions of NFeoB involved with *Vc*FeoC binding. Our findings thus reveal the first structure of *Vc*FeoC and how this small protein uses its truncated winged β-sheet to bind to *Vc*FeoB, lending further insight into the role of FeoC within the Feo system.

## Experimental methods

### Materials

The codon-optimized genes encoding *Vibrio cholerae* serotype O1 FeoC (*Vc*FeoC; Uniprot identifier C3LP26) and *Vibrio cholerae* serotype O1 (strain M66-2) FeoB (Uniprot identifier C3LP27) were synthesized by GenScript. Materials used for buffer preparation, protein expression, and protein purification were purchased from standard commercial suppliers and were used as received. Isotopically-enriched ammonium chloride (^15^NH_4_Cl) and glucose (globally ^13^C_6_-labeled) were purchased from Cambridge Isotope Laboratories and used as received. Detergents were purchased from Sigma-Aldrich, stored at −20 °C, and used as received. D_2_O was purchased from MilliporeSigma and used as received.

### Expression and purification of VcFeoC

The cloning, expression, and purification of *Vc*FeoC was similar to our previous FeoC preparations [17]. Briefly, DNA encoding the gene for *Vibrio cholerae* serotype O1 FeoC (*Vc*FeoC; Uniprot identifier C3LP26) with an N-terminal (His)_6_ tag, maltose binding protein followed by a Tobacco Etch Virus (TEV)-protease cleavage site (ENLYFQG) was sub-cloned into a pET45b(+) vector, transformed into chemically-competent BL21 (DE3) *E. coli* cells (Millipore Sigma, Burlington, MA), plated on Luria-Bertani (LB) agar plates containing 100 μg/mL ampicillin (final concentration), and incubated at 37 °C overnight. A single colony was used to generate large-scale (4× 1 L) cell cultures grown in LB supplemented with 100 μg/mL ampicillin. Cells were grown at 37 °C until reaching an OD_600_ of 0.6-0.8 at which point the cells were cold shocked briefly at 4 °C. For isotopically-enriched samples, a 100 mL LB starter culture treated with 100 μg/mL ampicillin was grown at 30 °C overnight and was used to inoculate 4× 1 L of M9 minimal medium containing ^15^NH_4_Cl and/or ^13^C_6_-glucose (Cambridge Isotope, Tewksbury, MA, USA). The cells were grown in this isotopically-enriched minimal media at 37 °C and shaken at 200 rpm until the OD_600_ reached 0.6-0.8 before a brief cold shock at 4 °C. Both natural abundance and isotopically-enriched samples were treated with isopropyl-β-D-1-thiogalactopyranoside (IPTG) to a final concentration of 1 mM and incubated at 18 °C with shaking at 200 rpm for 16-20 hours before harvesting by centrifugation at 4,800 ×*g*, 10 minutes, 4 °C. Cell pellets were resuspended in resuspension buffer (50 mM Tris, pH 7.5, 200 mM NaCl, 5% v/v glycerol), treated with approximately 50-100 mg solid phenylmethylsulfonyl fluoride (PMSF), and lysed by microfluidization (Microfluidics, Westwood, MA, USA). The lysate was clarified by ultracentrifugation at 163,000 ×*g* for 1 hour at 4 °C. The supernatant was applied to two tandem 5 mL MBPTrap HP columns (Cytiva, Marlborough, MA) and purified as previously described [17]. Fractions containing the target protein were concentrated using a 15 mL Amicon with 30 kDa molecular-weight cutoff (MWCO) spin concentrator, buffer exchanged into TEV-protease cleavage buffer (50 mM Tris, pH 8.0, 200 mM sodium chloride, 5% v/v glycerol, 1 mM TCEP, 0.5 mM ethylenediaminetetraacetic acid (EDTA)), and concentrated to 1 mL. The concentrated sample was treated with TEV protease and rocked at 4 °C overnight. The TEV-treated sample was purified by size-exclusion chromatography (SEC) using a 120 mL Superdex 75 column equilibrated with SEC buffer (25 mM Tris, pH 7.5, 100 mM sodium chloride, 5% v/v glycerol); cleaved, purified *Vc*FeoC was concentrated using a 15 mL 3 kDa MWCO spin concentrator. This purification approach yielded *ca*. 1-3 mg *Vc*FeoC L^−1^ of cell culture. Protein purity was assessed using 20% SDS-PAGE.

### NMR spectroscopy of VcFeoC

Each NMR sample contained *ca*. 2 mg of protein and was prepared in 50 mM sodium phosphate (pH 6.0), 5 mM sodium chloride with either 10% or 99% v/v D_2_O. Samples prepared in 99% D_2_O were exchanged from H_2_O using a PD-10 desalting column (Cytiva, Marlborough, MA). The PD-10 column was treated with 1.5 CVs of D_2_O, equilibrated with 1.5 CVs of NMR buffer prepared in D_2_O (50 mM sodium phosphate, pD 6.0, 5 mM NaCl), and eluted using 2 CVs of buffer. NMR datasets were acquired at 25 °C on a Bruker 600 MHz spectrometer equipped with a cryogenic probe. Heteronuclear single quantum coherence (HSQC) experiments were used to establish that *Vc*FeoC was folded and served as a basis for protein backbone assignments. Standard triple resonance experiments (CBCA(CO)NH, HNCACB, HNCO, and HN(CA)CO) were collected to assign the protein backbone [22–25]. A series of two-, three-, and four-dimensional nuclear Overhauser effect spectroscopy (NOESY) data were collected for combinations of natural abundance and isotopically-labeled (^15^N, ^13^C, and ^15^N/^13^C) protein samples. Protein dynamics were evaluated by ^1^H-^15^N heteronuclear nuclear Overhauser effect (XNOE) analysis. Data were processed with NMRPipe/nmrDraw or NMRFx and analyzed using NMRViewJ [26–29].

### Structural calculations

Structural calculations in torsion angle space were carried out using CYANA [30]. Upper interproton distance limits of 2.7, 3.3, and 5.0 Å were used for NOE cross-peaks of strong, medium, and weak intensities, respectively. Appropriate corrections of interproton distance limits were made for pseudoatoms. The TALOS+ Server was used to determine dihedral restraints that were incorporated into structural calculations based on amide proton, amide nitrogen, HͰ, CͰ, Cβ, and carbonyl carbon chemical shifts [31]. PyMOL was employed to prepare structural figures [32]. The atomic coordinates for *Vc*FeoC were deposited in the RCSB database (PDB ID 7U37). NMR chemical shifts and corresponding structure refinement parameters were deposited in the BMRB database (BMRB accession number 30995).

### Expression and purification of VcFeoB

The gene encoding for the *Vibrio cholerae* serotype O1 (strain M66-2) FeoB protein (Uniprot identifier C3LP27) was engineered to contain a C-terminal TEV-protease cleavage site (ENLYFQS) followed by a (His)_6_ tag for affinity chromatography purification and subcloned into the pET-21a(+) expression plasmid. This plasmid was transformed into chemically-competent BL21 (DE3) *E. coli* expression cells similar to the MBP-*Vc*FeoC construct. Large-scale expression of the protein was accomplished in 12 baffled flasks charged with 1 L of modified Terrific Broth supplemented with 100 μg/mL ampicillin. Growth was carried out at 37 °C with shaking at 200 rpm and monitored until an OD_600_ of 1.5-1.75 was reached. Flasks containing cells and media were then cold-shocked for at 4 °C before inducing protein expression with the addition of IPTG to a final concentration of 1 mM. Protein expression was carried out at 18 °C with shaking of 200 rpm overnight. After 18-20 h, the cells were harvested by centrifugation at 4800 ×*g*, 12 min, 4 °C. Cell pellets were resuspended in resuspension buffer (25 mM Tris, pH 7.5, 100 mM sucrose) and flash-frozen in N_2_ (*l*) before storage at −80 °C.

All purifications of *Vc*FeoB were carried out at 4 °C unless otherwise noted. Briefly, frozen cells containing the expressed protein were thawed and supplemented with solid PMSF (approximately 50-100 mg) before being lysed using a Q700 ultrasonic cell disruptor (QSonica, Newtown, CT) operating at 80% maximal amplitude, 30 s pulse on, 30 s pulse off, for a total duration of 12 min total pulse-on time. The lysate was then spun at 10000 ×*g* for 1 h to separate cellular debris and suspended membranes. The supernatant was decanted and ultracentrifuged at 163000 ×*g* for 1 h. Pelleted membranes were then washed, resuspended, and rehomogenized in resuspension buffer. Protein concentration was measured using the detergent-compatible (DC) Lowry assay (Bio-Rad Laboratories, Hercules, CA) before being flash-frozen on N_2_ (*l*) and stored at −80 °C. Membranes containing the *Vc*FeoB protein were thawed and solubilized with vigorous stirring for 3 h at 4 °C by the addition of a 10% (w/v) stock n-dodecyl-β-d-maltopyranoside (DDM) to a final concentration of 1% (w/v) detergent and 3-5 mg/mL total protein concentration. Insoluble material was then pelleted by ultracentrifugation at 163,000 ×*g* for 1 h before applying the supernatant to a 5 mL HisTrap HP column (Cytiva, Marlborough, MA) charged with Ni^2+^ and equilibrated with 10 column volumes (CVs) of wash buffer (25 mM Tris, pH 8, 100 mM sucrose, 300 mM NaCl, 1mM TCEP and 0.05 % (w/v) DDM) containing 21 mM of imidazole. After application of the clarified lysate, the column was then washed with 10 CVs of wash buffer containing 21 mM of imidazole. The protein was eluted by the wash buffer containing 150 mM imidazole. Fractions containing *Vc*FeoB were concentrated with a 15 mL Amicon 100 kDa MWCO spin concentrator (Millipore Sigma, Burlington, MA) and buffer exchanged into wash buffer using a PD-10 desalting column (Cytiva, Marlborough, MA). Protein purity was assessed via 15% SDS-PAGE analysis. Through this purification method, *ca*. 3-5 mg of pure *Vc*FeoB was obtained L^−1^ of cultured cells.

### Titration of VcFeoB into VcFeoC

Purified *Vc*FeoB and *Vc*FeoC samples were buffer exchanged into the *Vc*FeoB wash buffer (25 mM Tris, pH 8, 100 mM sucrose, 300 mM NaCl, 1mM TCEP and 0.05 % (w/v) DDM). HSQC spectra were collected as *Vc*FeoB was titrated into 100 μM *Vc*FeoC samples at stoichiometric (mole:mole) ratios of 0:1, 0.25:1, 0.5:1, and 1:1. Protein precipitation was observed at stoichiometric ratios greater than 1:1, preventing acquisition of further titration datapoints. The dissociation constant (K_d_) was estimated by integrating 1D ^1^H NMR spectra over the region in which *Vc*FeoC amide signals were detected (6.5-9.5 ppm) for all titration points. The fraction of *Vc*FeoB bound to *Vc*FeoC was determined by calculating the ratio of free *Vc*FeoC signal after *Vc*FeoB titration relative to unbound *Vc*FeoC, and these data were used to generate a binding isotherm. Data were processed using NMRFx [27], data integrations were carried out using dataChord Spectrum Analyst (One Moon Scientific, Inc., Westfield, NJ), and the binding isotherm fit was determined using Igor Pro (Wavemetrics, Lake Oswego, OR).

### Predicted docking model of VcNFeoB and VcFeoC

A homology model of *Vc*NFeoB was predicted by using ColabFold [33], which applies the AlphaFold2 structural prediction approach using MMseqs2 and HHSearch [34]. Docking studies were carried out for the lowest-energy predicted model in combination with the NMR structure of *Vc*FeoC by utilizing the ClusPro online server without modification to the default settings and restraints [35, 36]. The selected docking model chosen was that with the lowest balanced, weighted score that was also consistent with the NMR titration data.

## Results

### Expression, purification, cleavage, and isolation of VcFeoC

The expression, purification, cleavage, and isolation of untagged *Vc*FeoC from the MBP-tagged construct was carried out similarly to previously described approaches for *E. coli* and *K. pneumoniae* FeoC proteins (*Ec*FeoC and *Kp*FeoC, respectively) (Fig. S1) [17]. Specific deviations from the earlier reported methods included: (1) isolation of *Vc*FeoC from the tagged protein by carrying out the Tobacco Etch Virus (TEV)-protease cleavage reaction at 4 °C instead of room temperature to maintain solubility and (2) purification in the absence of reducing agent given that *Vc*FeoC lacks redox-sensitive Cys residues unlike *Ec* and *Kp*FeoC. This approach gave rise to yields of *ca*. 1-3 mg of highly pure protein L^−1^ of cell culture (Fig. S1). Notably, some reports suggest that the Feo system is only operative in an oligomerized, trimeric form; however, we see no evidence for formation of trimeric *Vc*FeoC, which behaves as a monomeric protein in solution based on data from gel filtration experiments (Fig. S1) and NMR analyses (*vide infra*). These results are consistent with previous results from our lab on NFeoAB (a fusion between FeoA and NFeoB), FeoA, and FeoC, which all appear as predominately monomeric species in our hands [17, 37, 38].

### Secondary structure of VcFeoC

Due to the small nature and good accumulation of recombinant *Vc*FeoC, nuclear magnetic resonance (NMR) spectroscopy was employed for structural and dynamics analyses. Gel filtration profiles under NMR conditions indicated that *Vc*FeoC was monodisperse and monomeric (Fig. S1B), consistent with previous findings reported for FeoC proteins [11, 17]. High-quality 2D ^1^H-^15^N heteronuclear single quantum coherence (HSQC) spectra were acquired for purified *Vc*FeoC (Fig. 1A), which demonstrated well-dispersed amide signals indicative of folded protein [39]. Although the ^1^H and ^15^N chemical shifts were generally insensitive to protein concentration (100-700 μM) and sample pH (6.0-8.0) (data not shown), backbone amide assignments were made at pH 6.0 due to optimal long-term protein stability and a decreased ^1^H-^2^H exchange of amide protons [40]. The established NMR conditions allowed for assignment of all backbone amide signals except for that of Val 61 (99% completion).

**Figure 1.**
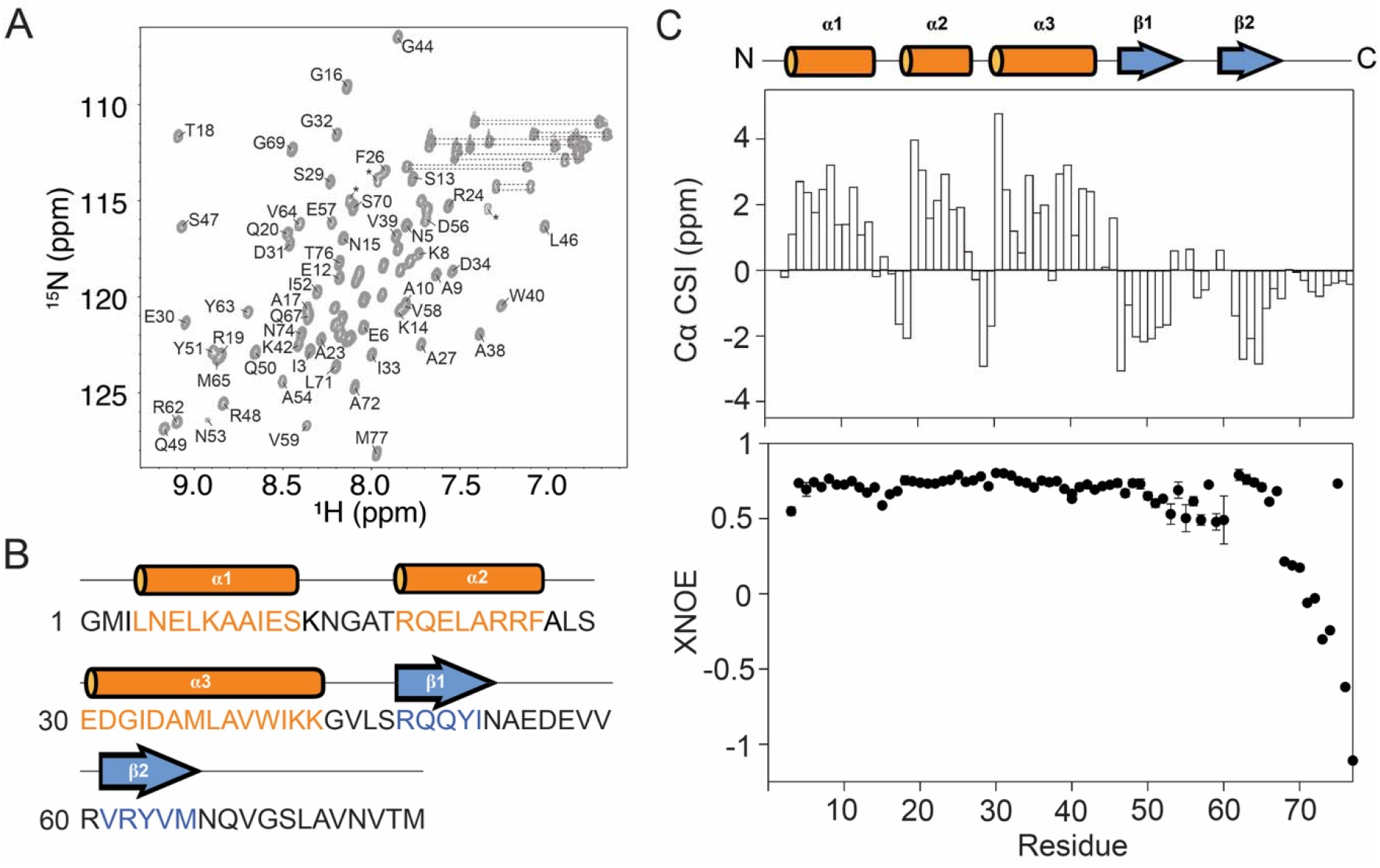
The secondary structure determination of *Vc*FeoC. **A**. Assigned ^1^H-^15^N HSQC NMR spectrum of 300 μM *Vc*FeoC at 298 K (50 mM sodium phosphate, pH 6.0, 5 mM NaCl, 10% v/v D_2_O). Assignments are generally explicit for residues in the less-crowded regions. Dashed lines represent signals corresponding to Asn and Gln side chains. Asterisks represent signals from Arg side chains. **B**. Amino acid sequence of *Vc*FeoC mapped with a cartoon of the corresponding secondary structure of each region. Residues belonging to α-helices are highlighted in orange (α labels) and include the following regions: Leu^4^ to Ser^13^ (α1), Arg^19^ to Phe^26^ (α2), and Glu^30^ to Lys^42^ (α3). The two-stranded β-sheet (β labels; blue) is composed of residues Arg^48^ to Ile^52^ (β1) and Val^61^ to Met^65^ (β2). Note: a single additional Gly residue at the N-terminus is present as a result of the TEV cleavage reaction. **C**. NMR chemical shift indices for backbone Cα atoms of *Vc*FeoC (top panel); positive values represent α-helical regions, negative stretches represent β-strand residues, and near zero values indicate unstructured and/or random coil regions. ^1^H-^15^N heteronuclear NOE (XNOE) data (bottom panel) indicate that *Vc*FeoC is largely structured with the exception of dynamic linkers (Lys^14^ to Thr^18^, Arg^25^ to Ser^29^, and Lys^43^ to Ser^47^), the dynamic wing region (Asn^53^ to Arg^60^), and the C-terminal tail (Asn^66^ to Met^77^). The amide signal of Val^75^ is overlaid with that of Glu^21^ (α2) and therefore gives an XNOE more consistent with a structured element. Error bars represent the standard

To determine the secondary structure of *Vc*FeoC, the Cα chemical shift indices (CSI) were analyzed based on assignment of triple resonance spectra (Fig. 1B-C) [41]. This analysis indicated that *Vc*FeoC is composed of three α-helices (α1, Leu^4^ to Ser^13^; α2, Arg^19^ to Phe^26^; α3, Glu^30^ to Lys^42^) and two β-strands (β1, Arg^48^ to Ile^52^; β2, Val^61^ to Met^65^), the latter of which are linked to form a short winged β hairpin terminating at an unstructured, dynamic C-terminal tail. Interestingly, negative Cα CSI values at residues immediately preceding the start of α2 (Thr^18^) and the start of α3 (Ser^29^) indicated the presence of N-terminal α-helix capping [42, 43], a structural feature that has been shown to impart additional stability to helices [44]. To characterize further the architecture of *Vc*FeoC, heteronuclear ^1^H-^15^N NOE (XNOE) data (Fig. 1C, bottom) were acquired. These data offered insight into the backbone dynamics and internal mobility for each signal, where XNOE measurements below *ca*. 0.8 are indicative of flexibility [45]. The XNOE analysis shows that *Vc*FeoC is largely structured except for three short linkers (Lys^14^ to Thr^18^, Arg^25^ to Ser^29^, and Lys^43^ to Ser^47^), the β hairpin wing residues (Asn^53^ to Arg^60^), and the C-terminus (Asn^66^ to Met^77^). Taken together, the XNOE findings both agree with the secondary structure determined from the Cα CSI data and are in general agreement with the structural architecture of previously studied FeoC proteins.

### Tertiary structure of VcFeoC

In order to determine the tertiary structure of *Vc*FeoC, ^15^N- and ^15^N/^13^C-isotopically enriched *Vc*FeoC samples were prepared, and 3D ^15^N-edited nuclear Overhauser effect (NOE), 4D ^15^N/^13^C-, and ^13^C/^13^C-edited NOE NMR spectra were acquired [46–49]. After data acquisition, structural calculations were carried out using a total of 648 interproton distance restraints derived from NOE data, 128 hydrogen bond restraints determined based on NOE crosspeak patterns, and 86 dihedral restraints based on backbone chemical shifts (Table 1). An ensemble of 20 refined structures with the lowest target function of 0.005 ± 0.002 Å^2^ was generated for *Vc*FeoC, and this ensemble exhibited good convergence based on root-meansquare deviations (RMSDs) of 0.12 ± 0.04 Å^2^ for backbone heavy atoms (Fig. 2; Table 1).

**Table 1.**
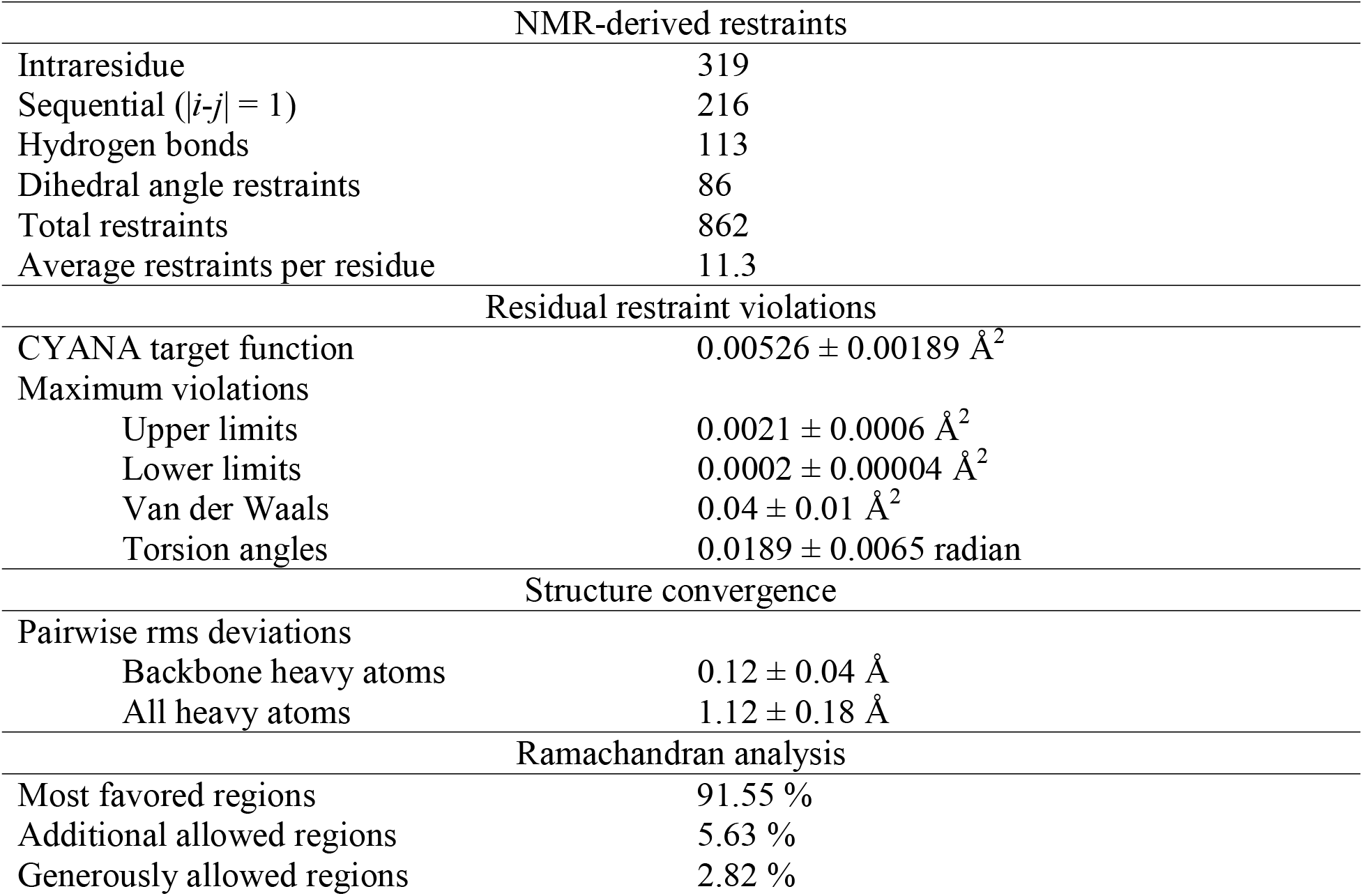
Structural restraints and refinement statistics for *Vc*FeoC.

**Figure 2.**
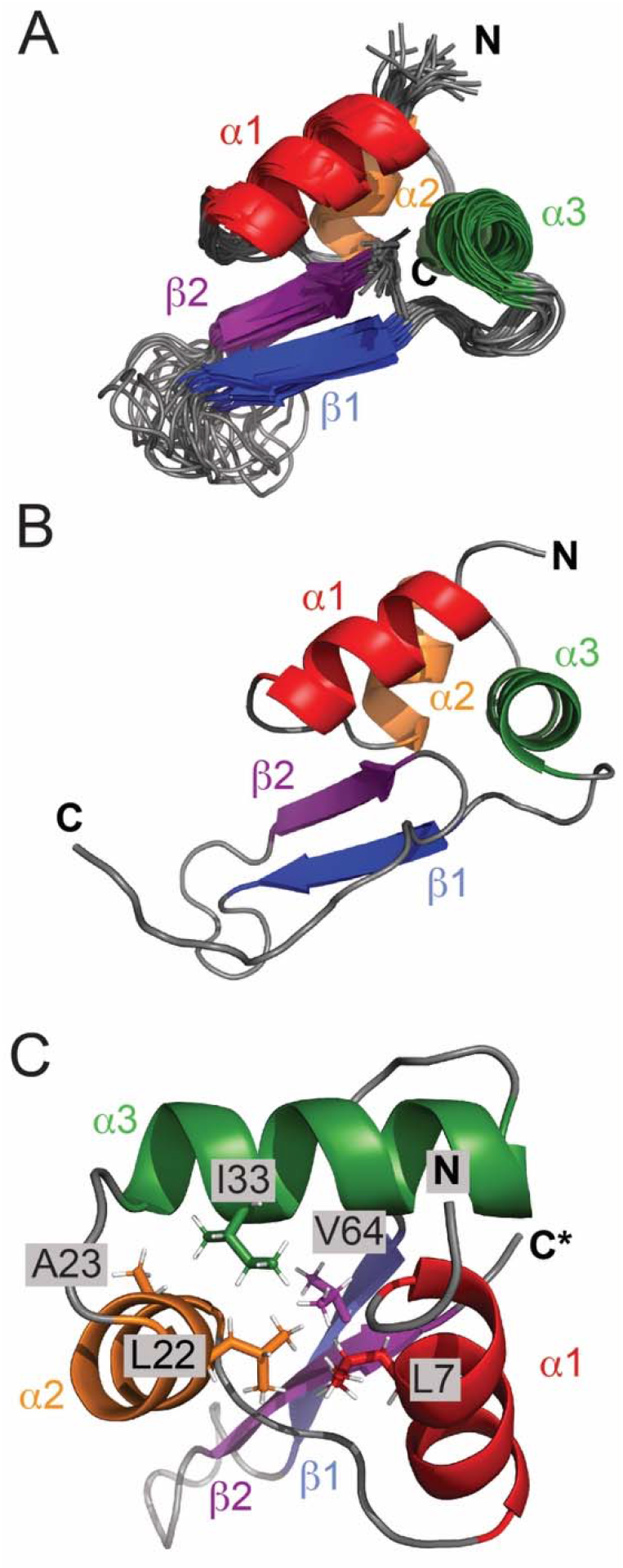
The NMR structure of *Vc*FeoC. **A**. Superposition of the 20 lowest-energy refined structures of *Vc*FeoC. The structured regions are Leu^4^ to Ser^13^ (α1; red), Arg^19^ to Phe^26^ (α2; orange), Glu^30^ to Lys^42^ (α3; green), Arg^48^ to Ile^52^ (β1; blue), and Val^61^ to Met^65^ (β2; purple). The ribbon representations include Gly 1 through Gln 67 to demonstrate the globular portion of the structure, but the dynamic C-terminal tail is truncated for clarity (represented by C*). **B**. Ribbon diagram of the full-length, lowest-energy target function structure of *Vc*FeoC. **C**. The hydrophobic core of *Vc*FeoC includes residues Leu^7^, Leu^22^, Ala^23^, Ile^33^, and Val^64^, which combined hold together the helix-turn-helix motif. The N- and C-termini are represented by ‘N’ and ‘C’ labels, respectively. Images in which the dynamic, unstructured C-terminus is truncated for figure clarity are labeled with ‘C*’.

As anticipated, the tertiary structure of *Vc*FeoC adopts a winged helix-turn-helix (HTH) structure featuring a three-helix bundle and a two-stranded antiparallel β-sheet that is connected by an unstructured wing. Long-range NOEs indicate that the hydrophobic core of *Vc*FeoC is composed of residues from α1-3 (Leu^7^, Leu^22^, Ala^23^, and Ile^33^) and β2 (Val^64^) (Fig. 2B-C). In order to quantify the similarity among *Vc*FeoC and other structurally characterized FeoCs, *Cα* RMSDs were determined of the following: *Kp*FeoC isolated from the X-ray crystal structure of *Kp*FeoC complexed with the N-terminal domain of *Kp*FeoB (*Kp*NFeoB) (1.793 Å; 29 Cαs) [10], apo *Kp*FeoC (1.770 Å; 22 Cαs) [11], and *Ec*FeoC (1.595 Å; 30 Cαs) (Table 2). Superpositioning of *Vc*FeoC upon these FeoC homologs highlights the similarity of the gross tertiary structure and, particularly, the conserved HTH domain (Fig. 3). Notably, the main structural differences among

**Table 2.** Cα RMSD s of the NMR-derived structure of *Vc*FeoC and other bacterial FeoC structures.

| Protein | PDB ID | RMSD (Å)<sup>a</sup> |
| --- | --- | --- |
| *Escherichia coli* FeoC | 1XN7 | 1.595 |
| *Klebsiella pneumoniae* FeoC | 2K02 | 1.770 |
| *Klebsiella pneumoniae* FeoC (FeoB-bound) | 4AWX | 1.793 |

**Figure 3.**
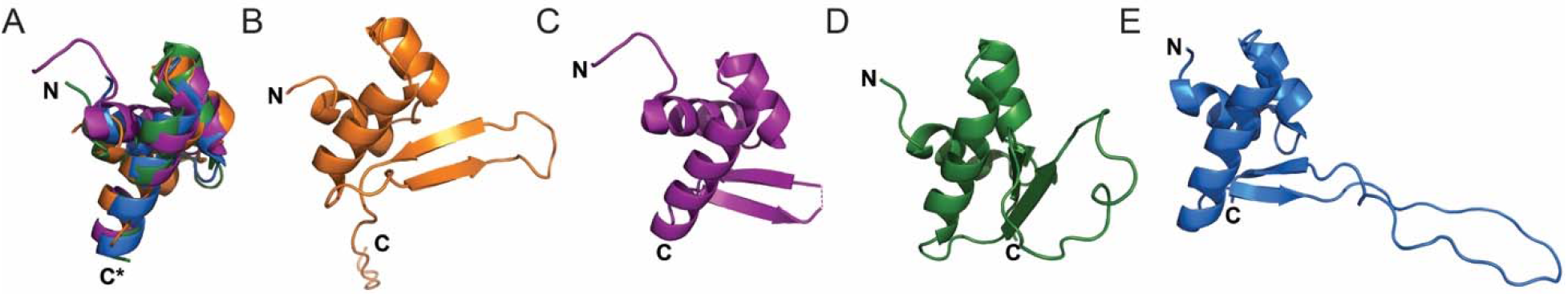
Comparisons of structurally-characterized FeoC proteins. **A**. Overlaid, truncated structures (residues 1-45) of the NMR-derived *Vc*FeoC (orange; PDB ID 7U37), the crystal structure of *Klebsiella pneumoniae* FeoC (*Kp*FeoC) isolated from the *Kp*FeoC-*Kp*NFeoB complex (purple; PDB ID 4AWX), apo *Kp*FeoC (green; PDB ID 2K02), and *Escherichia coli* FeoC (*Ec*FeoC, blue; PDB ID 1XN7). The superpositioning of these structures demonstrates the similarity within the conserved three-helix bundle (approximately residues 1-45 of each protein). **B-E**. Ribbon diagrams of the full-length NMR structure of *Vc*FeoC (**B**), crystallized *Kp*FeoC bound to NFeoB (**C**), the NMR structure of apo *Kp*FeoC **(D)**, and the NMR structure of apo *Ec*FeoC (**E**). These comparisons illustrate the heterogeneity observed within the winged regions of the winged-helix motif among FeoC proteins. The N- and C-termini are represented by ‘N’ and ‘C’ labels, respectively. Images in which the C-terminus is truncated for figure clarity are labeled with ‘C*’. ^a^ Root-mean-square deviation calculated between Cα atoms of matched residues

FeoCs are variations in the length of the β-strands, the extent of the β-sheet length, and the diversity in the length of the unstructured wing. Intriguingly, whereas both *Ec* and *Kp*FeoC have long, Cys-rich wings that serve to bind [4Fe-4S] clusters under anoxic conditions [16, 17], *Vc*FeoC features a shorter wing lacking Cys residues that cannot bind an [Fe-S] cluster. Despite this change, the wing of *Vc*FeoC is still quite dynamic, although it is incapable of sampling as much three-dimensional space as *Ec*- or *Kp*FeoC wings, due to the size differences. It is possible that this shorter wing region of *Vc*FeoC may be more constrained in space and may actually mimic the cluster-bound forms of *Ec*/*Kp*FeoC, which are known to be more compact [17].

### Interactions between VcFeoB and VcFeoC

Previous work by Hung and coworkers demonstrated the formation of a tight complex between *Kp*FeoC and the guanine dissociation inhibitor (GDI) domain of *Kp*NFeoB, suggesting a role for FeoC in the direct regulation of Fe^2+^ transport [10]. However, these studies were carried out with apo *Kp*FeoC, and the dynamic wing region was unresolved in the electron density; whether the cluster-bound form were capable of binding to *Kp*NFeoB was not explored. Moreover, these studies were unfortunately limited in that *Kp*FeoC was only tested for interactions with the soluble N-terminal domain of FeoB, not the intact membrane protein. In contrast, bacterial adenylate cyclase two-hybrid (BACTH) assays conducted by Weaver *et al*. indicated interaction of intact, full-length *Vc*FeoB and *Vc*FeoC under *in vivo* conditions [12]. Variant studies suggested that the interactions occurred between the N-terminal region of *Vc*FeoB and residues Glu^29^ and Met^35^ of *Vc*FeoC [12], but a direct observation of these interactions had not been determined.

Thus, we sought next to determine whether *Vc*FeoC binds to full-length *Vc*FeoB *in vitro* and, if so, to identify the binding interface of *Vc*FeoB-*Vc*FeoC, which could inform previous *in vivo* immunoprecipitation findings [20]. However, probing these interactions *in vitro* required the nontrivial preparation of large amounts of full-length *Vc*FeoB. After multiple optimization attempts, suitable heterologous expression of *Vc*FeoB featuring a C-terminal (His)_6_ tag was achieved. Solubilization in n-dodecyl-β-D-maltoside (DDM) and subsequent purification reproducibly resulted in the isolation of 2-3 mg of highly pure *Vc*FeoB L^−1^ of cells culture (Fig. S2). To determine whether *Vc*FeoC interacts with the intact *Vc*FeoB *in vitro*, 2D ^1^H-^15^N HSQC NMR spectra of *Vc*FeoC were acquired as *Vc*FeoB was titrated into various stoichiometric ratios (mole:mole) of *Vc*FeoC (Fig. 4). Given that the large size of *Vc*FeoB (*ca*. 85 kDa) exceeds the limit of detection of NMR, formation of the *Vc*FeoB-FeoC complex (*ca*. 94 kDa) results in a broadened and decreased NMR signal of the observed (unbound) *Vc*FeoC (*ca*. 9 kDa). Importantly, a control detergent-to-*Vc*FeoC titration was also performed to ensure that the decreased *Vc*FeoC signal intensity was the result of binding to *Vc*FeoB and not adventitious interactions with DDM micelles (Fig. S3). The overlaid HSQC data indicate that *Vc*FeoC binds *Vc*FeoB using the following regions on *Vc*FeoC: Asn^15^, Gly^16^, and Thr^18^ of the HTH domain; Glu^30^ of linker 2 (analogous to Glu^29^ identified in the previous BACTH studies [12]); Asp^34^ and Ala^38^ to Trp^40^ of α3; Leu^46^ and Ser^47^ of linker 3; Arg^48^ to Gln^50^ and Arg^62^ to Val^64^ β-sheet; and Glu^57^ of the wing (Fig. 4). We failed to observe any sign of interactions with the analogous Met^35^ in the previous BACTH studies. Attempts to titrate *Vc*FeoB into *Vc*FeoC beyond a 1:1 (mole:mole) ratio resulted in sample precipitation; however, the existing data still facilitated estimation of the binding affinity for complex formation. To do so, the 1D ^1^H signal over the amide region (6.5-9.5 ppm) was integrated, the fraction of *Vc*FeoC bound to *Vc*FeoB was determined for each titration point, and the binding isotherm for complex formation was evaluated (Fig. S4) [50, 51]. Using this analysis, we approximate that *Vc*FeoC binds to *Vc*FeoB with a modest K_d_ of *ca*. 25 μM, much weaker than the truncated apo *Kp*FeoC/*Kp*NFeoB complex.

**Figure 4.**
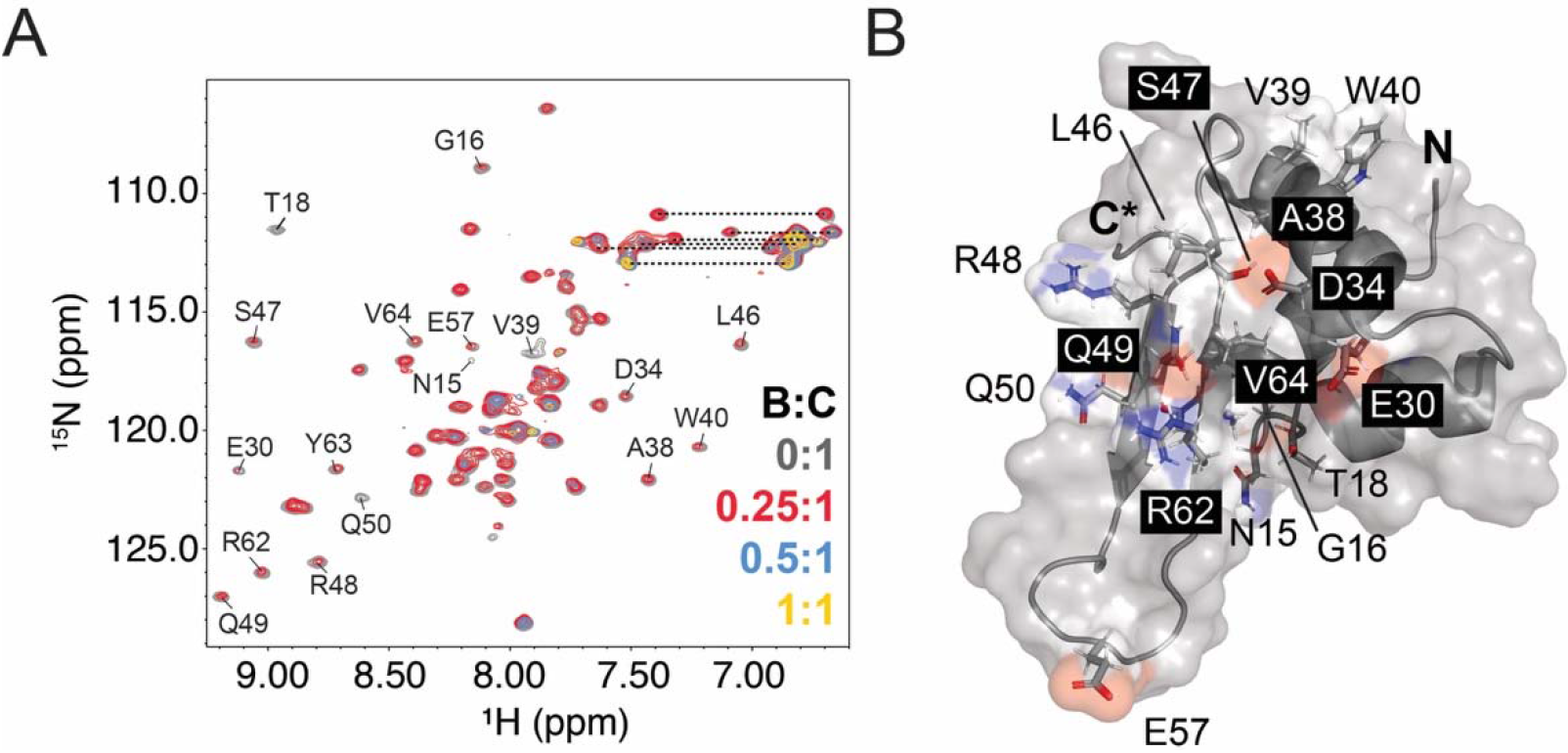
*Vc*FeoC binds to intact *Vc*FeoB in a 1:1 complex. **A**. Overlaid ^1^H-^15^N HSQC spectra of *Vc*FeoB titrated into 100 μM *Vc*FeoC (gray) at stoichiometric ratios (mole:mole) of 0.25:1 (red), 0.5:1 (blue), and 1:1 (yellow) at 298 K (50 mM Tris, pH 8.0, 100 mM sucrose, 200 mM NaCl, 0.05% dodecyl-β-D-maltoside, 1 mM TCEP). *Vc*FeoB (*ca*. 80 kDa) exceeds the limit of detection by NMR, and the broadened loss of *Vc*FeoC signals represents formation of the *Vc*FeoB-*Vc*FeoC complex. Labels are included and correspond to *Vc*FeoC signals that broaden the most rapidly due binding at the *Vc*FeoB/*Vc*FeoC interface. Dashed lines represent Asn and Gln side chain signals. **B**. A surface and cartoon representation of the truncated, lowest-energy structure of *Vc*FeoC (residues 1-67) indicating that residues within the helix-turn-helix (HTH) region (Asn^15^, Gly^16^, and Thr^18^), linker 2 (Glu^30^), α3 (Asp^34^, Ala^38^-Trp^40^), linker 3 (Leu^46^ and Ser^47^), β-sheet (Arg^48^-Gln^50^, Arg^62^-Val^64^), and wing (Glu^57^) broaden rapidly due to interaction with *Vc*FeoB. Labels are included for all residues for which broadened signals are observed except for Tyr^63^ that is on the opposite face of β2. The N- and truncated C-terminus is represented by ‘N’ and ‘C*’ labels, respectively.

As we presumed that *Vc*FeoC interacts with the N-terminal domain of *Vc*FeoB (*Vc*NFeoB) based on previous data [10], but our NMR titrations only give us a spectroscopic and structural handle for *Vc*FeoC, we then sought to understand the interaction between *Vc*FeoC and *Vc*NFeoB better through modeling approaches. As several structures of NFeoB homologs exist in the PDB, we determined a homology model of *Vc*NFeoB, and we used the lowest-energy model to dock *Vc*FeoC onto *Vc*NFeoB. Several docking models predicted interactions in a similar orientation to those shown in Fig. 5A, which demonstrates that electrostatic and hydrophobic interactions reinforce binding between Switch I/Switch II and the GDI domains of *Vc*NFeoB with the winged β-sheet of *Vc*FeoC, consistent with our NMR findings (*vide supra*).

**Figure 5.**
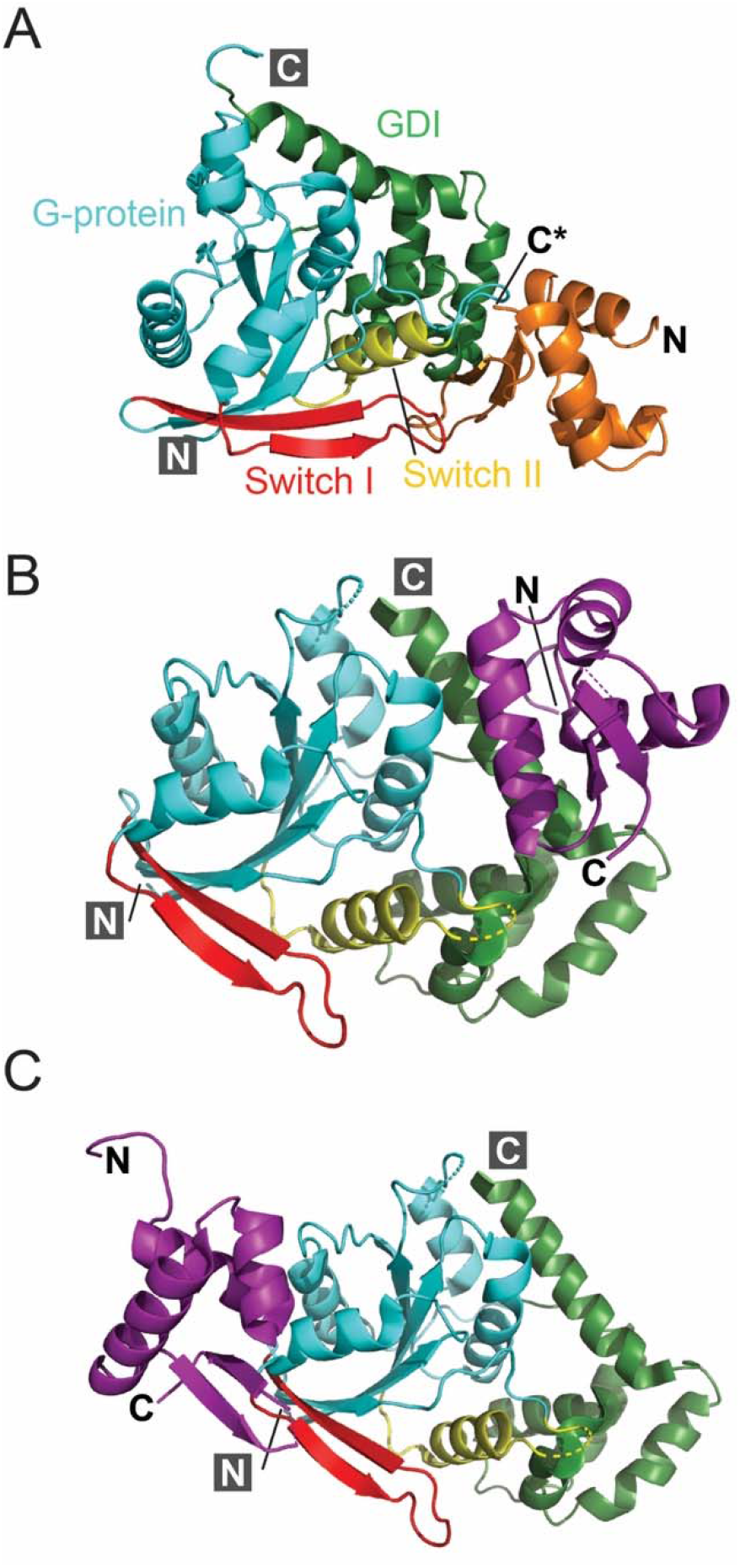
*Vc*FeoC-*Vc*NFeoB docking model and its comparison to the *Kp*FeoC-*Kp*NFeoB cocrystal structure. **A**. Docking studies of *Vc*FeoC (orange) and the homology model of *Vc*NFeoB suggest interactions of the *Vc*FeoC β-sheet and wing with Switch I (red), Switch II (yellow), and GDI (green) domains. **B**. Studies of *Kp*FeoC (purple) co-crystallized with *Kp*NFeoB (PDB ID 4AWX) indicate that *Kp*FeoC α3 interacts with NFeoB by means of hydrogen bonds, salt bridges, and hydrophobic interactions with the GDI (green) and Switch II (yellow) domains [10]. **C**. Extended crystal contacts throughout the crystalline lattice indicate an alternative mode of binding of *Kp*FeoC (purple) to *Kp*NFeoB in which the *Kp*FeoC wing interacts with the Switch I region of *Kp*NFeoB (red). Images in which the dynamic, unstructured C-terminus is truncated for figure clarity are labeled with ‘C*’. The G-protein domain is colored in cyan. The N- and C-termini are represented by ‘N’ and ‘C’ labels, respectively.

These results are similar to those observed in the *Kp*NFeoB/*Kp*FeoC complex X-ray crystal structure (PDB ID 4AWX; Fig 5B,C). Unfortunately, in that structure the asymmetric unit (ASU) of this complex was ambiguous and suggested that *Kp*FeoC could interact with *Kp*NFeoB via hydrogen bonding, electrostatic interactions, and interactions of hydrophobic residues between two different regions: the GDI domain on a single *Kp*NFeoB protomer and the Switch II region of the neighboring *Kp*NFeoB protomer (Fig. 5B) [10]. Our modeling data suggest that both could be operative, at least for *Vc*FeoC, which may represent a constitutive mimic of the holo, [Fe-S] cluster-bound form of FeoC, which was absent from the *K. pneumoniae* complex structure.

## Discussion

Although the function of FeoC remains disputed, it is clear that this poorly conserved protein serves a regulatory function that is important for Fe^2+^ transport in several γ-proteobacteria [12, 21, 52]. Many of these organisms are pathogenic prokaryotes, including notable problematic pathogens such as *Salmonella enterica* [52, 53], *V. cholerae* [12], and *K. pneumoniae* [10]. Our lab has demonstrated that the role of some FeoCs is likely dependent on the binding of an oxygen-sensitive [Fe-S] cluster binding in the dynamic wing regions of FeoC [17], contrasting earlier studies suggesting that these cluster-binding FeoCs could be oxygen-tolerant [16]. Studies of *S. enterica* FeoC further confirm that FeoC is oxygen-sensitive and could regulate FeoB levels under changing metabolic conditions [53]. Unfortunately, the oxygen-sensitive nature of the [Fe-S] cluster makes structural determination of cluster-replete FeoCs challenging [17]. However, some FeoC proteins in pathogens like *V. cholerae* lack the necessary cluster-binding residues, prohibiting [Fe-S] cluster binding, yet these proteins remain functionally important [12, 17, 20]. These observations have led us to hypothesize that the functional aspect of FeoC may either be located at a structural site outside of the [Fe-S] cluster-binding residues, or that FeoCs lacking [Fe-S] cluster binding could be constitutively active and always capable of affecting iron transport.

To this end, we determined the NMR structure of *Vc*FeoC, which is generally similar to the previously solved *Ec* and *Kp*FeoC structures (Fig. 3 and Table 2). As expected, *Vc*FeoC features the conserved N-terminal, trihelical HTH domain but differs at the C-terminal winged β-sheet [10, 11]. Two main differences are observed between *Vc*FeoC and its [Fe-S] cluster-binding homologs: *Vc*FeoC features a shorter β-sheet and wing regions and has a long, disordered C-terminal tail (Fig. 3) [11]. In particular, we believe that the observed differences in the winged β-sheet are due to differences in [Fe-S] cluster binding capabilities: *Ec*- and *Kp*FeoC bind [4Fe-4S] clusters requiring long, dynamic wings that undergo conformational changes to accommodate this cofactor [11, 16, 17], whereas *Vc*FeoC does not. We know that [Fe-S] cluster binding in *Ec*- and *Kp*FeoC results in compaction of structure [17], and we believe that *Vc*FeoC could naturally mimic this more compact structure without need of [Fe-S] cluster binding in order to affect function via protein-protein interactions.

FeoC is known to interact with other components of the Feo system, although a consensus on function still seems unclear. BACTH assays of the *V. cholerae* Feo system show that FeoB and FeoC interact [12], and immunoprecipitation studies demonstrate that FeoA, FeoB, and FeoC could form a complex, albeit very large [20]. Interestingly, this work proposed a requirement for FeoA but not FeoC in complex formation and suggested that FeoC could serve to regulate FeoB levels [20]. Moreover, the same assays implicated two *Vc*FeoC residues (Glu^29^ and Met^35^) in giving rise to interactions with FeoB, but other participating residues were not identified. Our structural work demonstrates that the winged β-sheet of *Vc*FeoC binds full-length *Vc*FeoB with a modest K_d_ (*ca*. 25 μM), and several residues including the previously proposed Glu^29^, but not Met^35^, are involved in *Vc*FeoB binding. Although the size of *Vc*FeoB prohibits the determination of the binding interface via NMR, docking studies suggest that the cavity formed by the GDI domain and the Switch I/II regions of *Vc*NFeoB acts as the binding receptacle for *Vc*FeoC. However, our data do not support the formation of a strong complex between only *Vc*FeoB and *Vc*FeoC, at least under these conditions. Considering the *in vivo* findings that FeoA, FeoB, and FeoC all interact in *V. cholerae* [20], it is plausible that the presence of *Vc*FeoA could strengthen the *Vc*FeoB-*Vc*FeoC interaction. In fact, this complex formation could even be nucleotide-mediated, and recent work from our lab has shown that FeoA-FeoB interactions can be facilitated by the presence of nucleotide [38]. Ultimately, these events may be related to Fe^2+^ translocation via the transmembrane domain, especially given FeoC’s interactions near the GDI domain that links directly to the first transmembrane helix. However, additional mechanistic and structural work is necessary to further probe this hypothesis, which is an exciting future avenue of research.

## Abbreviations

BACTH: bacterial adenylate cyclase two-hybrid
CV: column volume
DDM: n-dodecyl-β-D-maltopyranoside
DtxR: diphtheria toxin repressor
*Ec*FeoC: *Escherichia coli* FeoC
EDTA: ethylenediaminetetraacetic acid
Fe^2+^: ferrous iron
Fe^3+^: ferric iron
FNR: fumarate and nitrate reductase
GDI: GDP dissociation inhibitor
HSQC: heteronuclear single quantum coherence
IPTG: isopropyl-β-D-1-thiogalactopyranoside
K_d_: dissociation constant
*Kp*FeoC: *Klebsiella pneumoniae* FeoC
LTTR: LysR transcriptional regulator
MWCO: molecular-weight cutoff
NMR: nuclear magnetic resonance
NOE: nuclear Overhauser effect
NOESY: nuclear Overhauser effect spectroscopy
PMSF: phenylmethylsulfonyl fluoride
RMSD: root-mean-square deviation
SH3: SRC homology 3
SEC: size exclusion chromatography
TEV: tobacco etch virus
*Vc*FeoB: *Vibrio cholerae* FeoB
*Vc*FeoC: *Vibrio cholerae* FeoC
*Vc*NFeoB: *Vibrio cholerae* NFeoB
XNOE: heteronuclear nuclear Overhauser effect

## Acknowledgements

This work was supported by NSF CAREER grant 1844624 (A. T. S. and M. L.), NIH-NIGMS grant R35 GM133497 (A. T. S. and J. B. B.), and in part by NIH-NIGMS grant T32 GM066706 (M. L.). Sequence searches utilized both database and analysis functions of the Universal Protein Resource (UniProt) Knowledgebase and Reference Clusters (http://www.uniprot.org) and the National Center for Biotechnology Information (http://www.ncbi.nlm.nih.gov/). We thank Prof. Michael F. Summers (UMBC) for generous access to NMR facilities, reagents, and technical support.

## Statements and Declarations

### Competing interests

The authors declare no competing financial interest.

## For Table of Contents Use Only

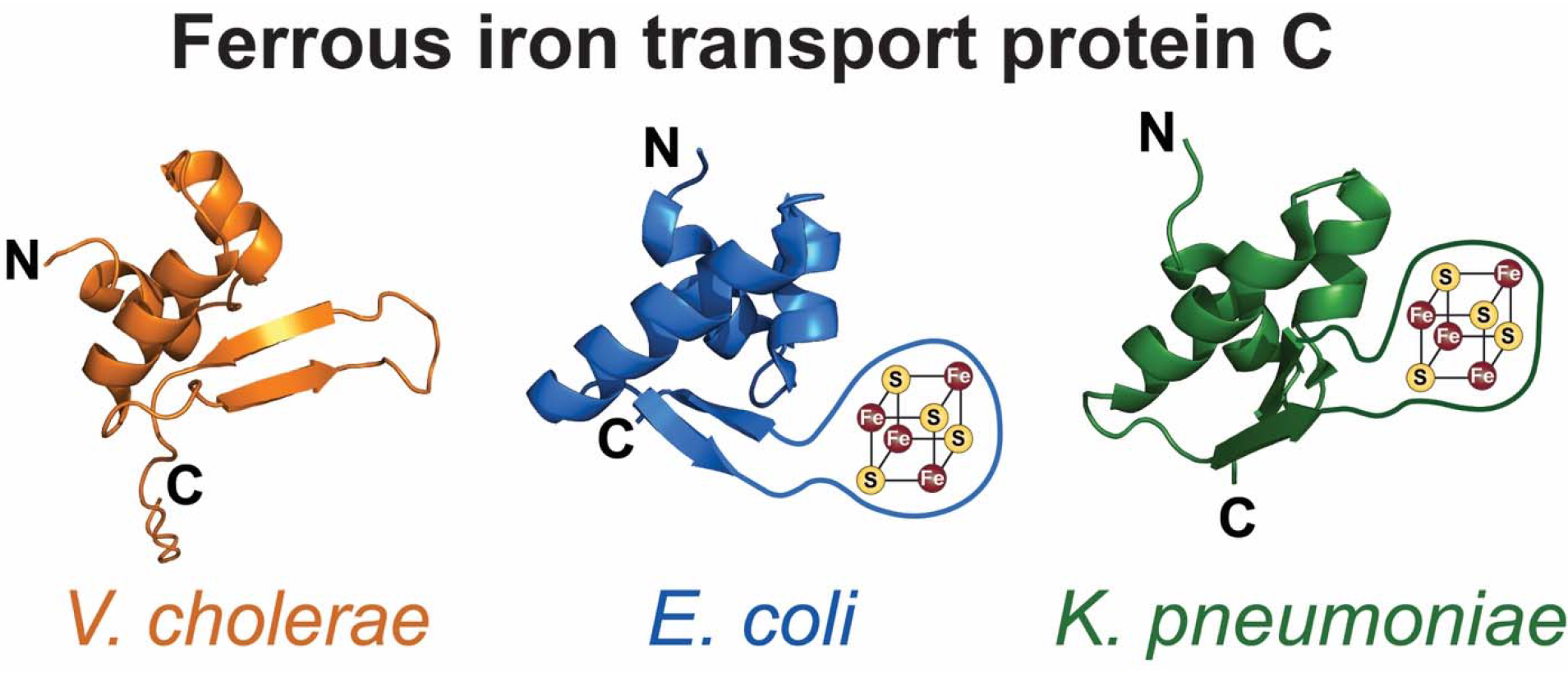

## Notes

### Competing Interest Statement

The authors have declared no competing interest.

