## Supplementary Information for "The structure of *Vibrio cholerae* FeoC reveals conservation of the helix-turn-helix motif but not the cluster-binding domain"

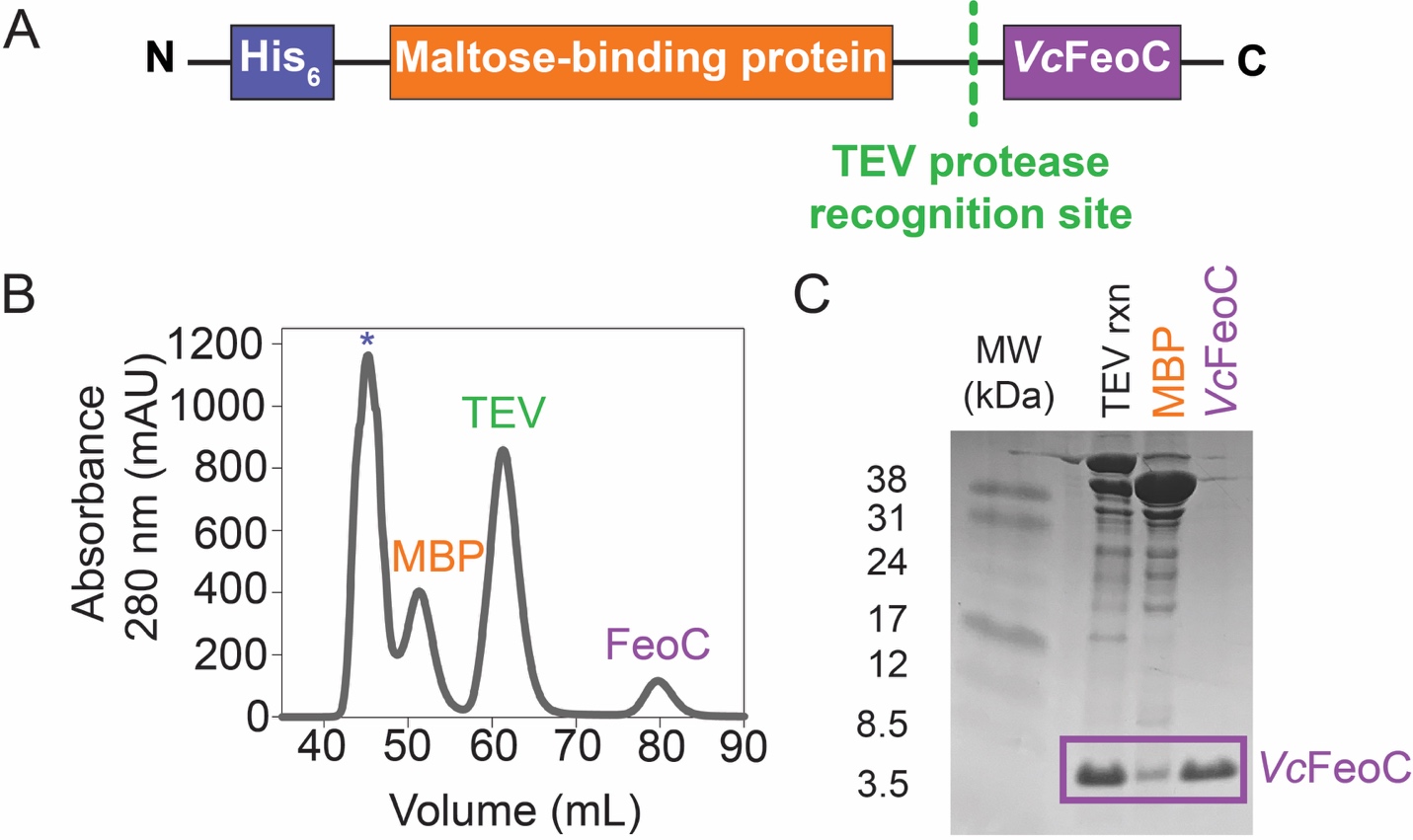

**Figure S1.** Construct design and purification of *Vc*FeoC. **A**. Cartoon representation of the MBP-*Vc*FeoC expression construct. An N-terminal (His)_6_ tag (blue box) is followed by maltose-binding protein (MBP) (orange box) and separated from *Vc*FeoC (purple box) by a Tobacco Etch Virus (TEV) protease recognition site (green). **B**. The Superdex 75 size-exclusion chromatography profile of MBP-*Vc*FeoC after TEV protease reaction shows good separation of aggregated protein (asterisk), cleaved MBP (orange), TEV protease (green), and *Vc*FeoC (purple). **C**. Representative 20% SDS-PAGE analysis highlighting the efficiency of the TEV cleavage reaction and the purity of *Vc*FeoC after size exclusion chromatography (final right-hand lane).

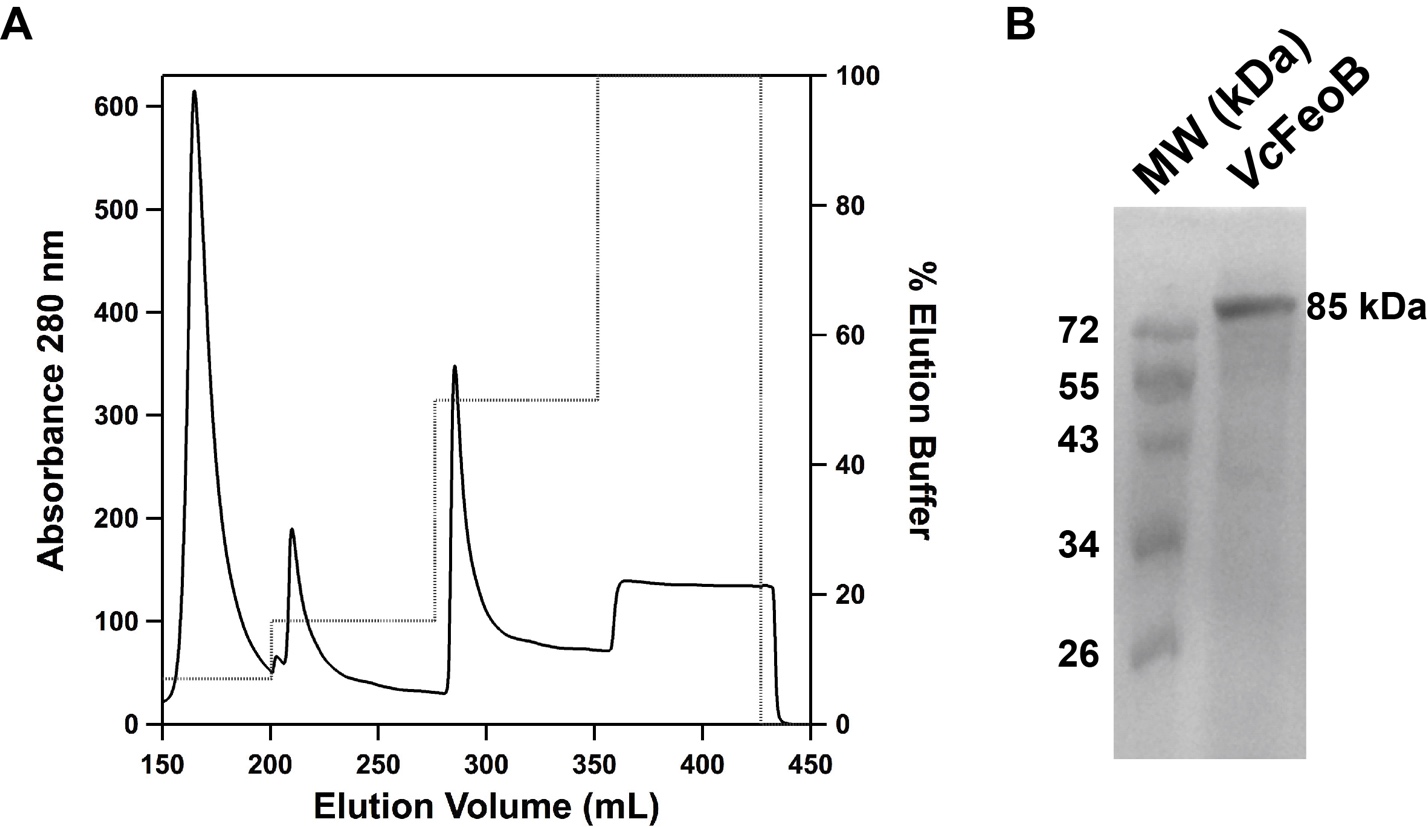

**Figure S2.** Immobilized metal affinity chromatography (IMAC) purification of *Vibrio cholerae* FeoB (*Vc*FeoB) solubilized in n-dodecyl-β-D-maltoside (DDM). **A**. IMAC chromatogram of the *Vc*FeoB purification demonstrates that the protein elutes at 50% elution buffer (50 mM Tris, pH 8.0, 100 mM sucrose, 200 mM NaCl, 0.05% DDM, 1 mM TCEP, 150 mM imidazole). **B**. 15% SDS-PAGE analysis demonstrating the purity of *Vc*FeoB after IMAC purification (right lane). The molecular weight of *Vc*FeoB is estimated at 85 kDa.

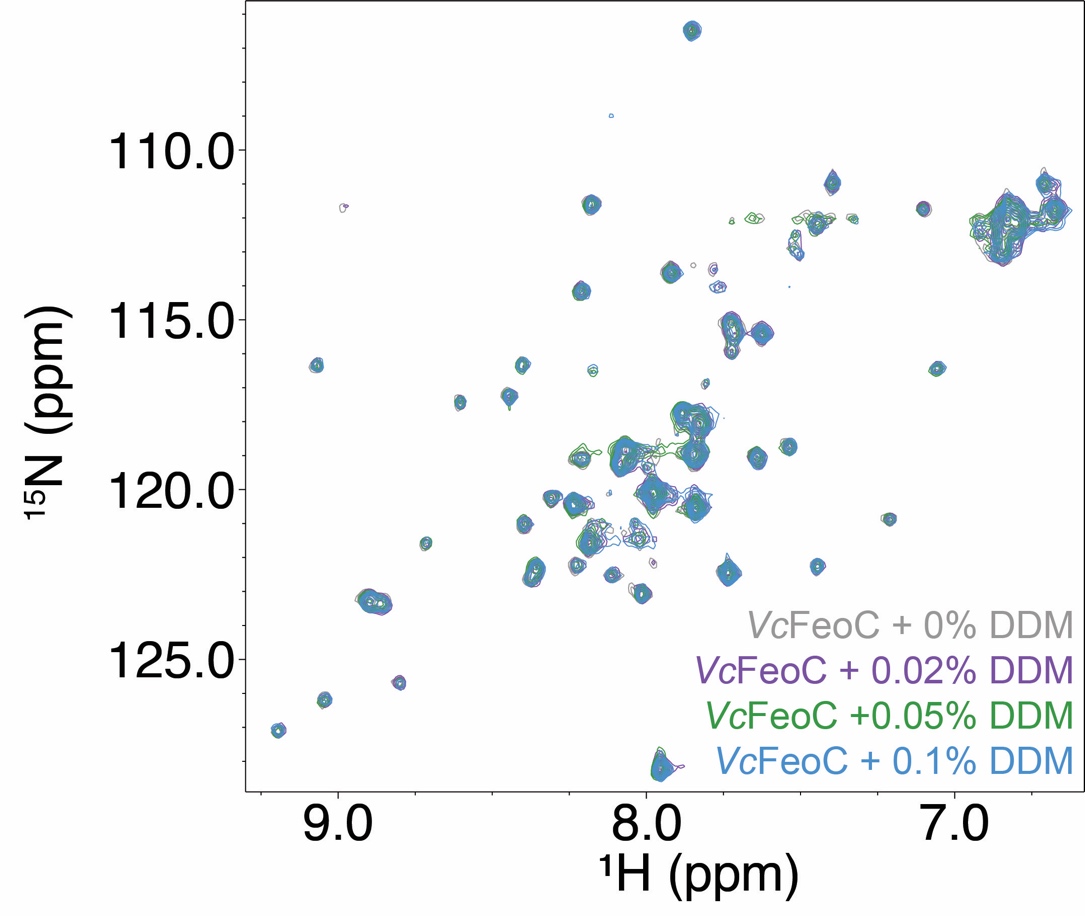

**Figure S3.** Overlaid ^1^H-^15^N HSQC spectra of *Vc*FeoC in the presence of increasing concentrations of n-dodecyl-β-D-maltoside (DDM). The spectra of *Vc*FeoC were acquired in the absence (0%) (gray), and the presence of 0.02% (purple), 0.05% (green), 0.1% (blue) w:v DDM, respectively. These results demonstrate that *Vc*FeoC does not interact adventitiously with DDM under the conditions employed for FeoB binding assays (50 mM Tris, pH 8.0, 100 mM sucrose, 200 mM NaCl, 1 mM TCEP, 0.05% DDM).

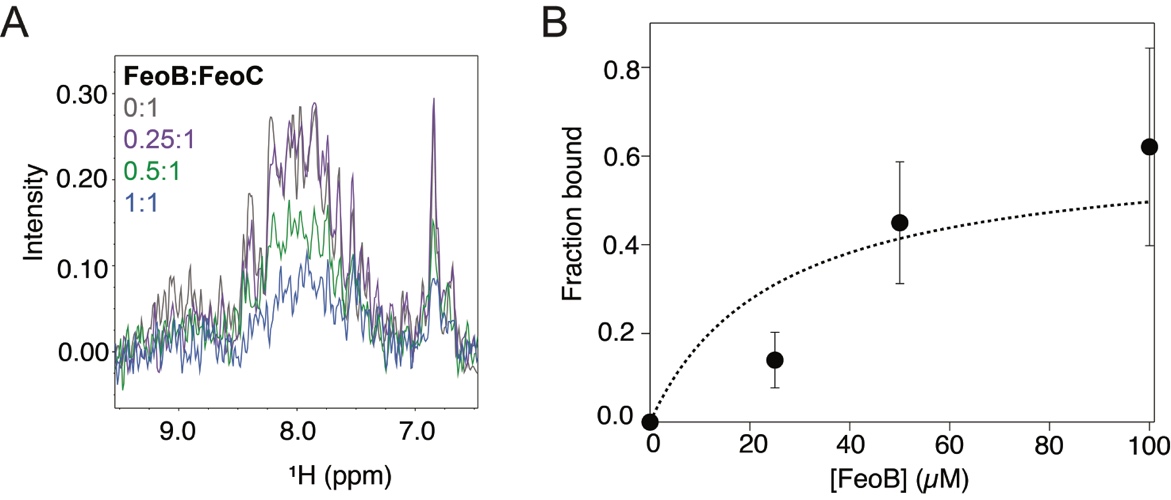

**Figure S4.** NMR titrations approximate the *Vc*FeoC-*Vc*FeoB binding affinity. **A**. The 1D ^1^H data extracted from 2D HSQC analyses were integrated over the region in which *Vc*FeoC amide signals are detected (6.5-9.5 ppm) at stoichiometric mole ratios (mol:mol) of FeoB to FeoC including 0:1 (gray), 0.25:1 (purple), 0.5:1 (green), and 1:1 (blue), respectively. Signal broadening and intensity loss is attributed to formation of the *Vc*FeoC-*Vc*FeoB complex (*ca*. 94 kDa) that exceeds the detection limit of NMR. **B**. The free-*Vc*FeoC signal (0:1, gray) was used to calculate the mole ratio of *Vc*FeoC bound to *Vc*FeoB as the membrane protein was titrated into *Vc*FeoC. The binding isotherm gives rise to a K_d_ of approximately 25 µM.
